# Probabilistic spike propagation shapes sympathetic output in mouse preganglionic neurons

**DOI:** 10.64898/2026.05.20.726575

**Authors:** Mallika Halder, Alan J Sokoloff, Yaqing Li, Michael Sawchuk, Brittney Ward, Shawn Hochman

## Abstract

Sympathetic preganglionic neurons (SPNs) provide the final pathway through which the central nervous system regulates autonomic function. SPN axons projecting to paravertebral sympathetic chain ganglia branch extensively and diverge across multiple segments, enabling amplification of central sympathetic commands through extensive postganglionic neuronal populations. Spike propagation along these projections has generally been assumed to occur reliably. However, most SPN axons are extremely small unmyelinated fibers, a structural feature predicted to reduce the safety factor for spike propagation.

Using an isolated mouse thoracic sympathetic chain preparation, we combined anatomical tracing with multi-site compound action potential recordings to assess conduction across SPN axons. Neurobiotin labeling revealed widespread rostrocaudal divergence through interganglionic nerves, while axon measurements confirmed that most SPN axons are small unmyelinated fibers. Across preparations, supramaximal recruitment of SPNs revealed substantial intertrial variability in compound responses, indicating frequent conduction failures. Failures were most prominent in slow-conducting axons and occurred in both branching interganglionic pathways and the unbranching axons within the splanchnic nerve. During repetitive activation, frequency dependent depression was observed at 1, 5 and 10Hz, but only slow-conducting branching axons exhibited pronounced depression.

Overall, these findings indicate that spike propagation in SPN axons may operate probabilistically rather than deterministically, with reliability strongly dependent on axonal subtype and recent activity history. We conclude that axonal conduction variability constitutes an intrinsic and dynamically regulated mechanism that shapes sympathetic output. By varying the recruitment of postganglionic populations, unreliable spike propagation in SPN axons introduces a previously unrecognized presynaptic gain-control mechanism, operating independently of central spike generation to modulate sympathetic output.

**SIGNIFICANCE:** Sympathetic preganglionic neurons provide the final pathway through which the central nervous system controls end-organs. These neurons project through the sympathetic chain where their axons branch extensively to recruit more numerous paravertebral postganglionic neurons. Spike propagation along these projections has generally been assumed to occur reliably. Here we show that this assumption is incorrect. Using anatomical tracing and electrophysiological recordings in mouse sympathetic chain preparations, we demonstrate that spike conduction in sympathetic preganglionic axons is frequently variable and prone to failure, particularly in the slowest-conducting unmyelinated fibers. Conduction variability was preferentially enhanced in branching axonal pathways during repetitive activation. These findings reveal that axonal conduction reliability represents an important presynaptic mechanism regulating the magnitude and variability of sympathetic output.

## INTRODUCTION

Axons are traditionally viewed as reliable conduits that faithfully transmit action potentials from neuronal somata to their synaptic targets. Variability in spike conduction can arise from branching architecture, axonal geometry, and activity-dependent changes in membrane excitability, allowing axons themselves to influence how information is transmitted within circuits (Goldstein and Rall, 1974; Levy, 1980). Increasing evidence indicates that axons are dynamic computational elements capable of modifying information transfer through neuromodulation, short-term plasticity, and branch-point filtering (Debanne, 2004; Debanne et al., 2011). While such mechanisms have been examined in central circuits, their contribution to autonomic signaling is uncertain.

Sympathetic preganglionic neurons (SPNs), located in the thoracic and rostral lumbar spinal cord, provide the final integrated pathway through which the central nervous system controls sympathetic output. SPNs integrate signals from central circuits and relay them to postganglionic neurons in the paravertebral sympathetic chain and prevertebral ganglia. SPNs comprise multiple distinct subpopulations (Alkaslasi et al., 2021; Blum et al., 2021). Those projecting to thoracic chain ganglia participate in the regulation of diverse physiological processes including cardiovascular control, metabolism, thermoregulation, and stress responses (Janig, 2022). SPN axons exit the spinal cord through segmental ventral roots and enter the paravertebral sympathetic chain via the white ramus. They often branch extensively as they course rostrocaudally through interganglionic nerves, forming divergent multisegmental projections across several ganglia (**Fig. 1A**)(Lichtman et al., 1980; Forehand et al., 1994). Through this anatomical organization, individual SPNs can influence many postganglionic neurons to amplify central commands onto much larger postganglionic populations, with estimated ratios of ∼200:1 in humans and 14:1 in mice (Purves et al., 1986). The presence of axonal branch points introduces geometric discontinuities that can reduce spike amplitude and conduction velocity or cause propagation failures. Although branch point conduction failure is well recognized in other neural systems (Goldstein and Rall, 1974; Levy, 1980; Debanne, 2004; Hari et al., 2022), it has not been widely considered as a factor influencing sympathetic output.

**Figure 1.**
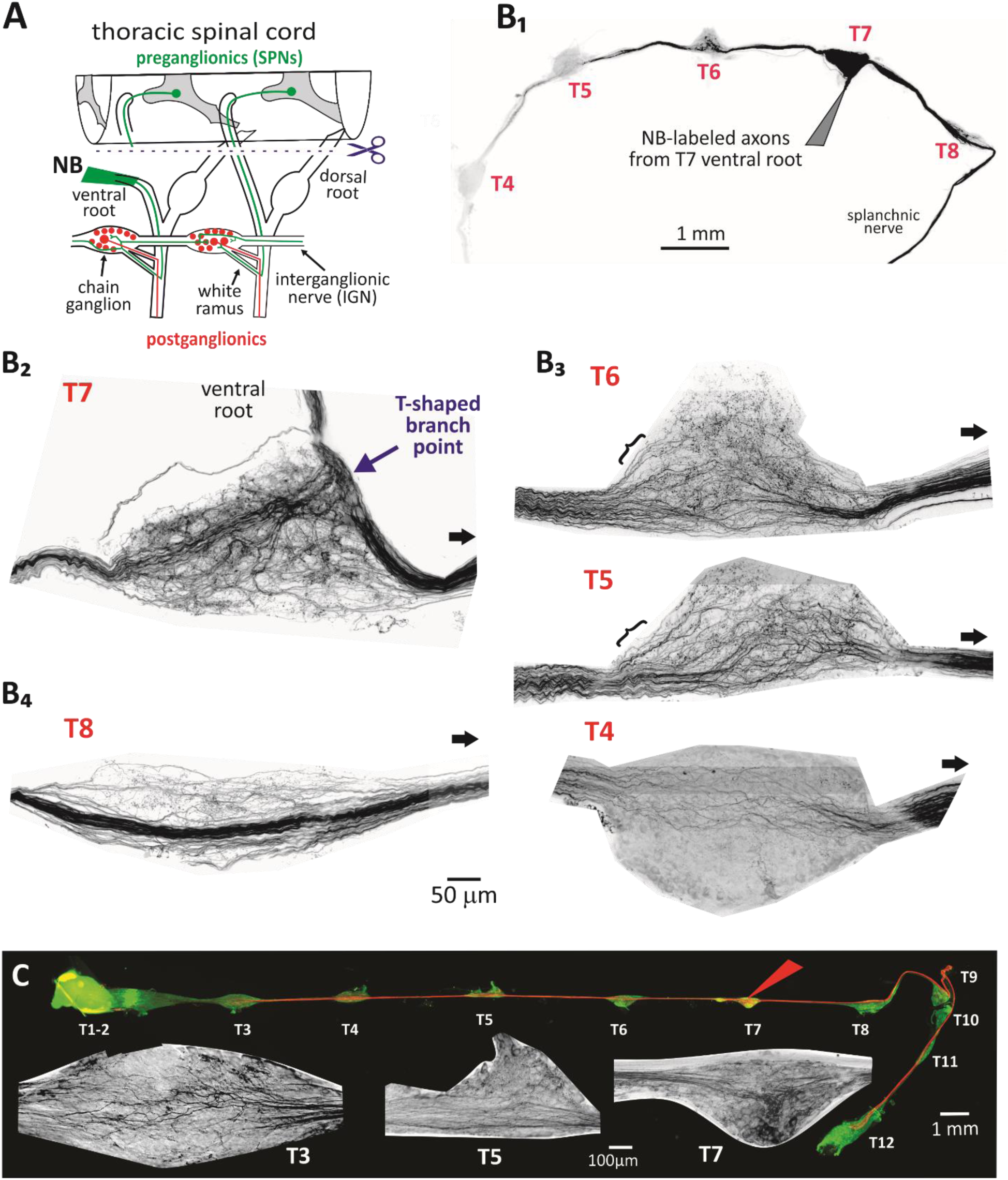
Thoracic SPN axons originating at the T7 spinal segment have broad multi-segmental divergent projections with multiple branching patterns. T7 SPN axons were labeled with Neurobiotin (NB) after diffusion from the T7 ventral root (n=2). **[A]** Anatomical overview. After cutting spinal roots, the spinal cord is removed from the vertebral canal and NB-containing solution is attached to a ventral root for anterograde labeling of segmental spinal SPN axons. **[B_1_]** NB^+^ labeling is seen along T3-T8 chain ganglia and in the splanchnic nerve [T9-12 ganglia are missing]. **[B_2_]** T7 SPN axons form a ‘T’ shaped branch point junction (arrow). **[B_3_]** Many T6 and T5 axons issue collaterals to form putative synapses before projecting to further rostral sites (see brackets at T5 and T4). Far fewer axons remain at T4. **[B_4_]** Most SPN axons that travel caudally are unbranching and may contribute to the splanchnic nerve which predominantly innervates the celiac ganglion. In panels B_2-4_, caudal direction is shown by a black arrow. **[C]** In another mouse, divergent rostral- and caudal-going NB^+^ labeling (red) can be seen along the length of the thoracic chain ganglia containing TH^+^ postganglionic neurons (green). T3, T5 and T7 ganglia insets are shown at magnified view (grayscale inverted and contrast enhanced) to show branching patterns.

Instead, spike conduction along SPN axons is generally assumed to remain reliable as signals propagate through the sympathetic chain (McLachlan, 2003). The structural features of SPN axons challenge this assumption. Nearly all mouse SPN axons are extremely small unmyelinated fibers with mean diameters near 0.4 µm and branch diameters that may approach ∼0.1 µm (Lewis and Burton, 1977). Such small axons are predicted to have high axial resistance and reduced safety factors for spike propagation, making them susceptible to conduction slowing, activity-dependent hyperpolarization, and conduction block. Small-diameter unmyelinated axons are also particularly sensitive to physiological mechanisms that regulates spike propagation, including activity-dependent hyperpolarization mediated by the α3 Na⁺/K⁺ pump, which acts as a metabolic excitability rheostat linked to recent firing history (Dobretsov and Stimers, 2005; Zhang and Sillar, 2012). These anatomical and physiological features suggest that spike conduction in SPN axons may be more variable than previously appreciated. If so, axonal conduction could represent an additional presynaptic mechanism shaping sympathetic output, complementing synaptic and network-level mechanisms.

Using an isolated mouse preparation with intact ventral roots and sympathetic chain ganglia (Halder et al., 2021), we test whether spike conduction in SPN axons is reliably maintained across their divergent projections in the sympathetic chain. Anatomical tracing examined SPN axon diameter, myelination, and branching architecture, while electrophysiological recordings characterized SPN axonal conduction properties. To determine whether conduction failures arise from axonal branching or from the small unmyelinated nature of SPN axons, we compared paravertebral responses recorded from branching interganglionic nerves, with those recorded from the adjacent splanchnic nerve with unbranching SPN axons projecting to prevertebral ganglia.

We demonstrate that conduction failures occur in a significant fraction of the SPN population. Slow-conducting axons exhibit substantial variability, with branching axons being particularly sensitive to frequency-dependent depression. That SPN axonal conduction variability can modulate the magnitude and temporal structure of sympathetic output, includes sympathetic preganglionic axons within a broader class of neural systems in which axons actively shape information transmission rather than serving as passive conduits.

## METHODS

### Anatomical studies

#### Neurobiotin labeling to assess preganglionic divergence

Details for surgical isolation of the intact thoracic sympathetic chain are provided in our earlier publication (Halder et al., 2021). Briefly, adult (8 weeks+) C57Bl/6 mice of both sexes were anesthetized with intraperitoneal injection of urethane (2g/kg). The thoracic vertebral column and adjacent ribs were excised and transferred to a dish containing ice cold, oxygenated (95% O_2_ / 5% CO_2_) high-Mg^2+^/low-Ca^2+^ extracellular fluid solution containing (in mM), [NaCl 128, KCl 1.9, MgSO_4_ 6.5, CaCl_2_ 1.1, KH_2_PO_4_ 1.2, glucose 10, NaHCO_3_ 26]. ACSF pH was adjusted to 7.4 after saturation with gas (95%O_2_, 5%CO_2_) at room temperature. Following a complete laminectomy and vertebrectomy, the spinal cord (SC) and dorsal roots (DR) were exposed and removed. The remaining thoracic chain ganglia, in continuity with communicating rami, spinal nerves and ventral roots were cleaned of excess fat and muscle. The tissue was transferred to a Sylgard 170 (Dow Inc.) silicone-bottomed chamber superfused with oxygenated extracellular solution (in mM), [NaCl 128, KCl 1.9, MgSO_4_ 1.3, CaCl_2_ 2.4, KH_2_PO_4_ 1.2, glucose 10, NaHCO_3_ 26], at ^∼^40ml/minute flow rate at 22C. To obtain evidence of widespread SPN divergence, we labelled SPN axons in the T7 or T10 ventral roots with Neurobiotin dissolved in distilled water (Vector Labs, 6%, RRID:SP110). This was loaded into an appropriately tapered glass capillary tube to enable root suction of the cut-end of either the T10 or T7 ventral root and left undisturbed for 6 hours at 22 °C. Subsequently, the preparation was removed from the perfusion chamber and fixed in 4% paraformaldehyde (0.5M phosphate, 4% paraformaldehyde, NaOH) for 2 hours. After fixation, the paravertebral thoracic chain was immersed in 20% sucrose solution and stored at 4°C. The sympathetic chain was subsequently isolated from stellate (T1 and T2) to T12/13. The dissected chain was washed overnight in PBS with Triton (PBS-T) solution containing 20% DMSO and 0.3% Triton-X. It was then placed in sealed vials filled with the same PBS-T solution and incubated in a 37°C water bath overnight. To prevent any reaction of DMSO with oxygen, we minimized air exposure in the vials. A foam float ensured the vial was completely submerged in the water bath. This step was repeated using a second PBS-T solution (20% DMSO, 0.1% Tween20, 0.1% Sodium Deoxycholate, 0.1% Tergitol, 0.3% Triton X 80%) with the same precautions regarding air exposure. Next, the tissue was transferred to a third PBS-T solution (20% DMSO, 0.3 M Glycine, 0.3% Triton X 80%) for three hours at 37°C in the water bath. Following this, an overnight PBS-T wash and 3-4 subsequent hourly washes were performed to fully remove any DMSO residue. The chain was subsequently incubated for 5-7 days with ExtrAvidin Cy3 (Sigma, 1:000, RRID:E4142). The tissue was then washed in PBS-T (5 x 1hr), then 50mM Tris-HCl (2 x 1hr followed by 1 x overnight) then again in 50mM Tris-HCl (2 hours) before mounting on prepared glass slides (four dots of nail polish were applied to the slides to slightly elevate the cover slip).

#### Assessment of axon counts and size

The T7 white ramus and the interganglionic pathway rostral and caudal to all paravertebral ganglia with Neurobiotin reaction were imaged for Neurobiotin immunoflouresence on a Nikon Crest Spinning Disk Confocal (60x objective, NA 1.49, optical resolution 232 nm, 0.108 microns/pixels X-Y, 0.150-micron Z-steps). Image stacks were deconvolved (Microvolution) and analyzed in FIJI for manual axon counts (Tubeness, Reslice) and cross-section measurement (Stardist, Analyze Particles) of Neurobiotin^+^ axons. We analyzed 120 optical slices from each rostral-caudal tract for 3 interganglionic nerve (IGN) regions (T5-T4, T-6-T5, and T7-T6) to assess axon count, diameter, and number. The data from these slices were averaged to obtain a measurement for each segment. This analysis was conducted using FIJI software. To evaluate differences in diameter and cross-sectional area (CSA) between IGN segments, we employed one-way ANOVAs followed by Dunn’s pairwise multiple comparison procedures. Values are reported as mean ± standard error (SEM).

#### Myelin basic protein and cholinergic axon dual immunolabeling

Following perfusion fixation as described above, the paravertebral chain with attached roots was stained with rat anti-myelin basic protein in a ChAT-mhChR2-eYFP (JAX:014546) mouse (Sigma, 1:50, MAB386) and chicken anti-GFP (Abcam, 1:100, AB3970). Whole mounts were washed overnight in PBS-T then incubated in primary antibodies for 3 days followed by wash in PBS-T (3 x 30 min). Tissue was incubated for 3 hours in a secondary antibody solution: Cy3 donkey anti-rat (Jackson Immunoresearch, 1:250, RRID:712-165-153) and FITC anti-chicken (Jackson Immunoresearch, 1:100, RRID:703-095-155). The tissue underwent a final wash in PBS-T (1 x 20 min) and 50mM Tris-HCl (2 x 20 min) before mounting on glass slides.

### Electrophysiological studies

#### Multisegmental *ex vivo* paravertebral preparation

Details are provided in our earlier publication (Halder et al., 2021). The electrophysiological component of presented experiments were undertaken in adult choline acetyltransferase (ChAT)-channelrhodopsin (ChR2) mice; driver line is ChAT-IRES-cre (JAX: 006410); reporter line is R26-ChR2-eYFP (JAX: 012569). Here we report predominantly on electrically evoked responses in relation to assessment of conduction failures. The chain isolation, extracellular solutions, and perfusion are as described above for Neurobiotin labeling. We focused on actions at room temperature as prior physiologic studies on multisegmental SPN actions on paravertebral postganglionic were undertaken at room temperature - 22°C (Blackman and Purves, 1969; Nja and Purves, 1977; Lichtman et al., 1980). The ganglia of thoracic paravertebral chain were made visible and glass suction electrodes were placed delicately to ensure stable recordings for extended periods. Recording electrodes were placed on the caudal cut ends of IGNs and splanchnic nerve with internal diameters ranging from 25-50 *µ*m (Halder et al., 2021). Glass suction electrodes were also positioned on thoracic ventral roots for stimulation (200-250 *µ*m tip diameter). All recorded data were digitized at 50 kHz (Digidata 1322A 16 Bit DAQ, Molecular Devices, U.S.A.) with pClamp acquisition software (v. 10.7 Molecular Devices). Recorded signals were amplified (5000x) and low pass filtered at 3 kHz using in-house amplifiers. As recordings of population axonal spiking (CAPs) could include synaptically-mediated postganglionic neuronal spiking responses, we blocked synaptic activity with the nicotinic acetylcholine receptor antagonist hexamethonium (100 μM) (Mason, 1962) and pancuronium (20 µM) to prevent possible intercostal muscle activity (both from Sigma-Aldrich).

#### Recruitment profiles for SPN volleys

We compared SPN axonal recruitment at multiple stimulus strengths and duration using electrical stimuli on ventral roots while recording evoked responses in the IGN caudal and in the splanchnic nerve. While recruitment profiles captured evoked responses from both anodic and cathodic stimuli, anodic stimuli commonly provided the largest responses and was used in most experiments. Electrical stimulation threshold was found to be ∼50µA, 50µs in the T12 IGN and splanchnic nerve recordings in most preparations with maximal recruitment typically achieved at 200µA, 200µs stimulation (n=5). To ensure supramaximal recruitment of slow-conducting axons, electrical stimulation intensity for experiments was kept at 200µA, 500µs, as longer pulse durations ensure recruitment of unmyelinated axons (Thompson et al., 1990; Brocker and Grill, 2013).

#### Quantification and analysis of evoked SPN CAPs

Because individual unmyelinated axons could not be uniquely resolved electrophysiologically, conduction failures were assessed as variability in population-evoked compound action potentials following supramaximal recruitment of SPN axons. Recorded CAPs were divided into conduction velocity (CV)-subpopulations depending on latency of arrival from time of stimulation. For each CV population, 10 episodes were rectified and integrated using Clampfit software (Molecular Devices). Background noise was removed from the rectified integral by subtracting the rectified integral obtained from an equal recorded duration prior to stimulation to provide a singular, comprehensive measure of CAP size. All 10 episodes were then averaged to ensure a more accurate representation of the CAP. Values are reported as mean percent of baseline ± standard error (SEM). A Student’s t-test was employed to determine statistical significance in comparing conditions. One-way repeated measures ANOVA for tests across multiple groups. For post-hoc detailed pairwise comparisons, a Bonferroni t-test analysis was utilized.

## RESULTS

### Anatomical assessment of CNS preganglionic (SPN) axon composition and projections

#### SPN projections issue complex multi-segmental branching patterns into thoracic chain ganglia

We utilized Neurobiotin intra-axonal labeling from ventral roots to understand the anatomical structure and projections of axons from mid to caudal thoracic spinal segments. While there was variability in distance associated with successful fills, several examples showed considerable rostrocaudal branching projections (**Fig. 1A, B_1_**). Notably, many SPNs issued diverging rostrocaudal collaterals after exiting the T7 white ramus, seen as T-shaped branch points (**Fig 1B_2_**) with additional branching projections seen across many rostrocaudal segments within the paravertebral sympathetic ganglia (**Fig. 1B_3,4_**). Counts were made of axons entering and exiting across two rostral and one caudal ganglia segment in the sympathetic chain shown in Fig 1B. We counted Neurobiotin^+^ axon numbers at sites of entry and exit from ganglia. Of the 165 SPN axons labeled in the T7 ramus, 125 axons projected rostrally into the T7-T6 interganglionic nerve (IGN), while 175 axons projected caudally into the T7-T8 IGN, indicating that most T7 axons bifurcated for rostrocaudal projections at this point (**Fig. 2A**). As shown, rostral axonal projections included those that bypass the ganglia without entering and those that enter the ganglia to issue putative synapses. At least some of these axons entering the ganglia reentered the IGN to project rostrally, consistent with electrophysiologically recorded divergent synaptic actions across multiple ganglia (Lichtman et al., 1980) (e.g. **Fig. 1B_3_**).

**Figure 2.**
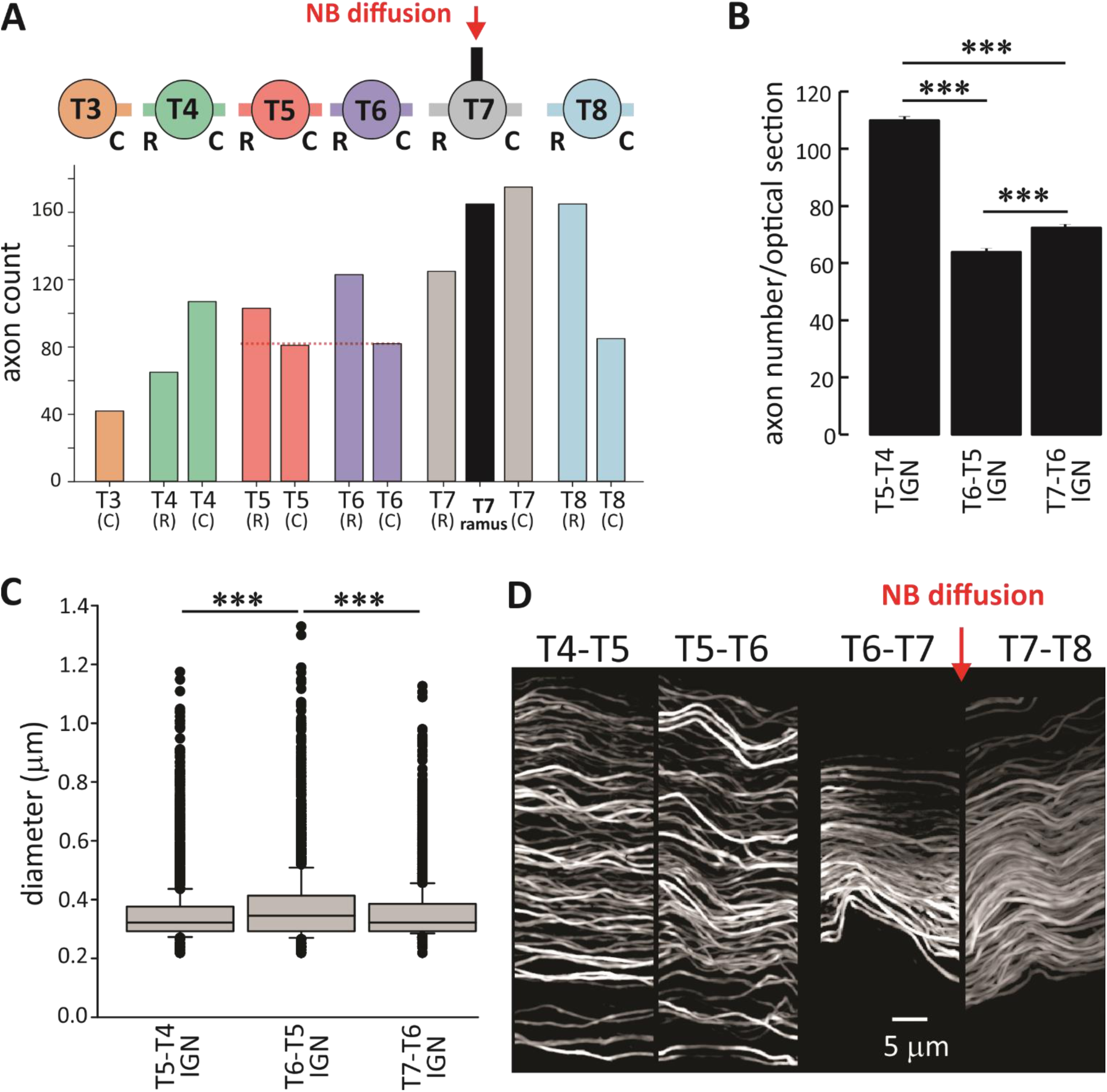
Assessment of axon anatomical properties along the interganglionic nerve. **[A]** Counts of Neurobiotin labeled axon numbers in the T7 ramus and in interganglionic nerves projecting rostrally (R) and caudally (C) between T3 and T8 sympathetic ganglia (counts are with longitudinal analysis of Z-stacks after deconvolution and tubeness at 0.25). Schematic above bar graph summarizes the anatomical basis behind presented counts. Note that as axons travel rostrally from T7, more axons appear to exit both T6 and T5 ganglia than enter (see dotted horizontal lines in red). Experiment schematic showing NB diffusion at the T7 ventral root. For all graphs, cross sections of the IGN between ganglia (red) were measured for axon count, mean cross sectional area and mean diameter of axons. **[B]** Increase in number of axons projecting across the T6 and T5 ganglia supports more axonal branching (red dotted line). Mean axon numbers in cross section counted across >100 transverse optical slices in each IGN are as follows: T7_R_-T6_C_ (73.2 ± 0.3), T6_R_-T5_C_ (64.4 ± 0.07), T5_R_-T4_C_ (110 ± 2.0) (***; p<0.001). Total axon numbers counted across sections are T7_R_-T6_C_ (8,646), T6_R_-T5_C_ (7,730), and T5_R_-T4_C_ (10, 835). **[C]** Box plot of axon diameter distributions across sections. Diameters were comparable between T7-T6 (0.355 ± 0.001µm), T6-T5 (0.372 ± 0.001µm) and T5-T4 (0.349 ± 0.001µm). One-way ANOVA identified statistical differences (***; p<0.001) with posthoc differences between T6-T5 and the other interganglionic regions as shown (p<0.001). **[D]** Axon trajectories in the IGN. Note that IGN axon locations closer to the white ramus of origin are more tightly clustered while axons at more locations spread out, presumably interlacing with axons from other segments.

**Figure 3.**
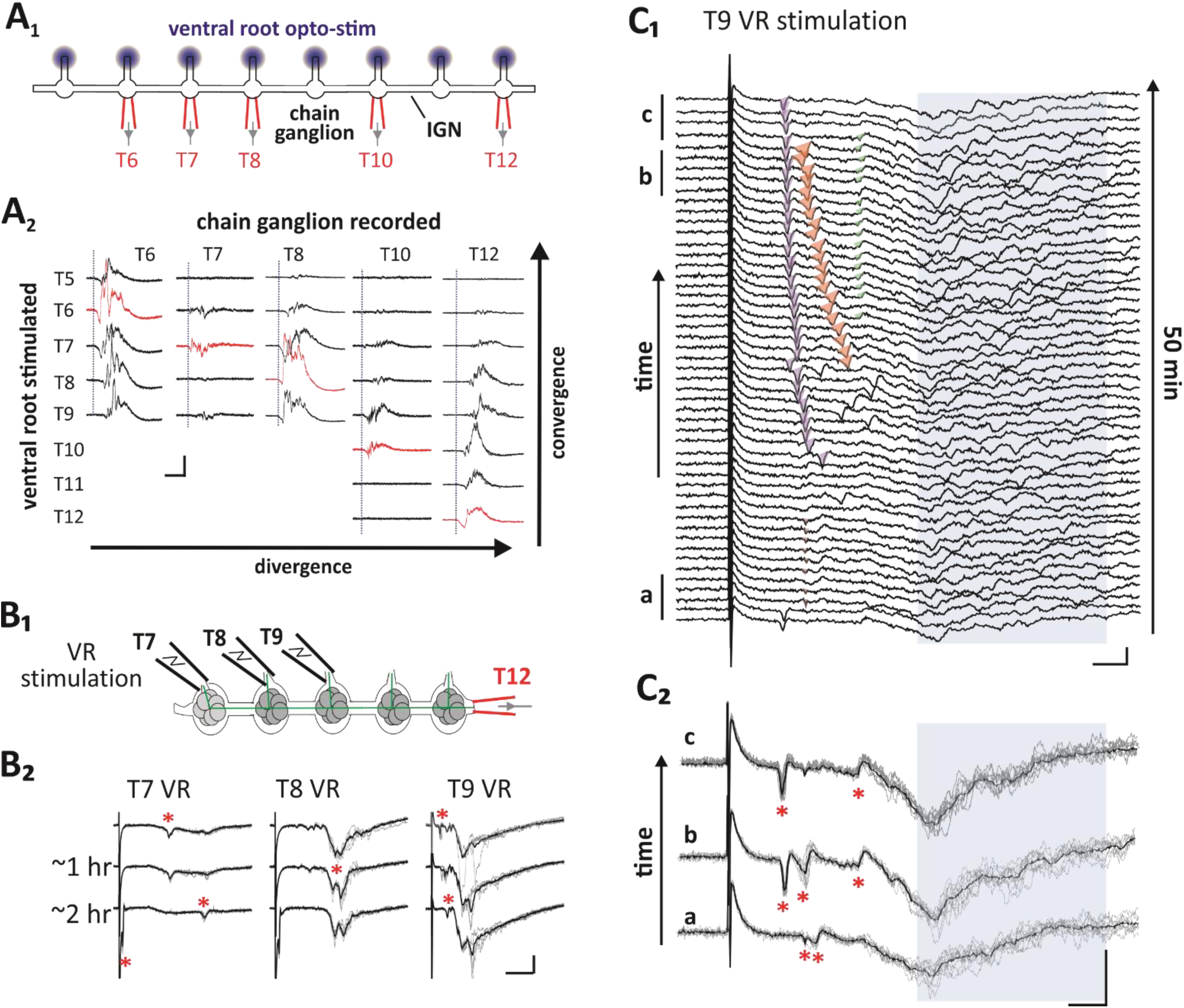
Multisegmental convergence, divergence and dynamic fluctuations in axonal recruitment over time. **[A_1_]** Experimental design showing supramaximal optical stimulation of SPNs in mid to lower thoracic ventral roots of ChAT-CHR2 mouse (represented by blue dots) with *en passant* ganglia recordings from listed ganglia using glass suction electrodes (illustrated in red), capturing the direct recruitment of SPN and postganglionic responses (no hexamethonium applied). **[A_2_]** Convergent and divergent projections are displayed as evoked responses from three episodes overlaid. The traces in red indicate responses in ganglia following stimulation of the same segment. Similar results were obtained from 4 mice. **[B_1_]** Experimental setup and the recording of CAPs from converging SPN axons at T11 IGN, originating from T7, T8, and T9 ventral roots. **[B_2_]** Changes in conduction were observed over a two-hour period. Shown is 5 episodes with average response overlaid in black. Note that evoked responses are larger when the stimulation site is closer to the recording site. Asterisks identify individual units undergoing time-dependent changes in recruitment. **[C_1_]** Shown is a raster plot of individual CAP records following T9 VR stimulation at one-minute intervals for 50 minutes (episodes are ordered chronologically from bottom to top). The emergence of two larger early units with progressively earlier latency (orange and purple shading) is consistent with increased conduction security. Other emergent or lost events are shaded in pink or green. The high variability in expression and arrival time in the later arriving unmyelinated axons (blue shaded region) prevents assessment of individual events necessitating characterization as population changes. **[C_2_]** Overlaid events in the 5-minute epochs identified in raster above further highlights changes in axonal recruitment. Scale bars: [A & B] 100μV, 20ms; [C] 100μV, 5ms.

**Figure 4.**
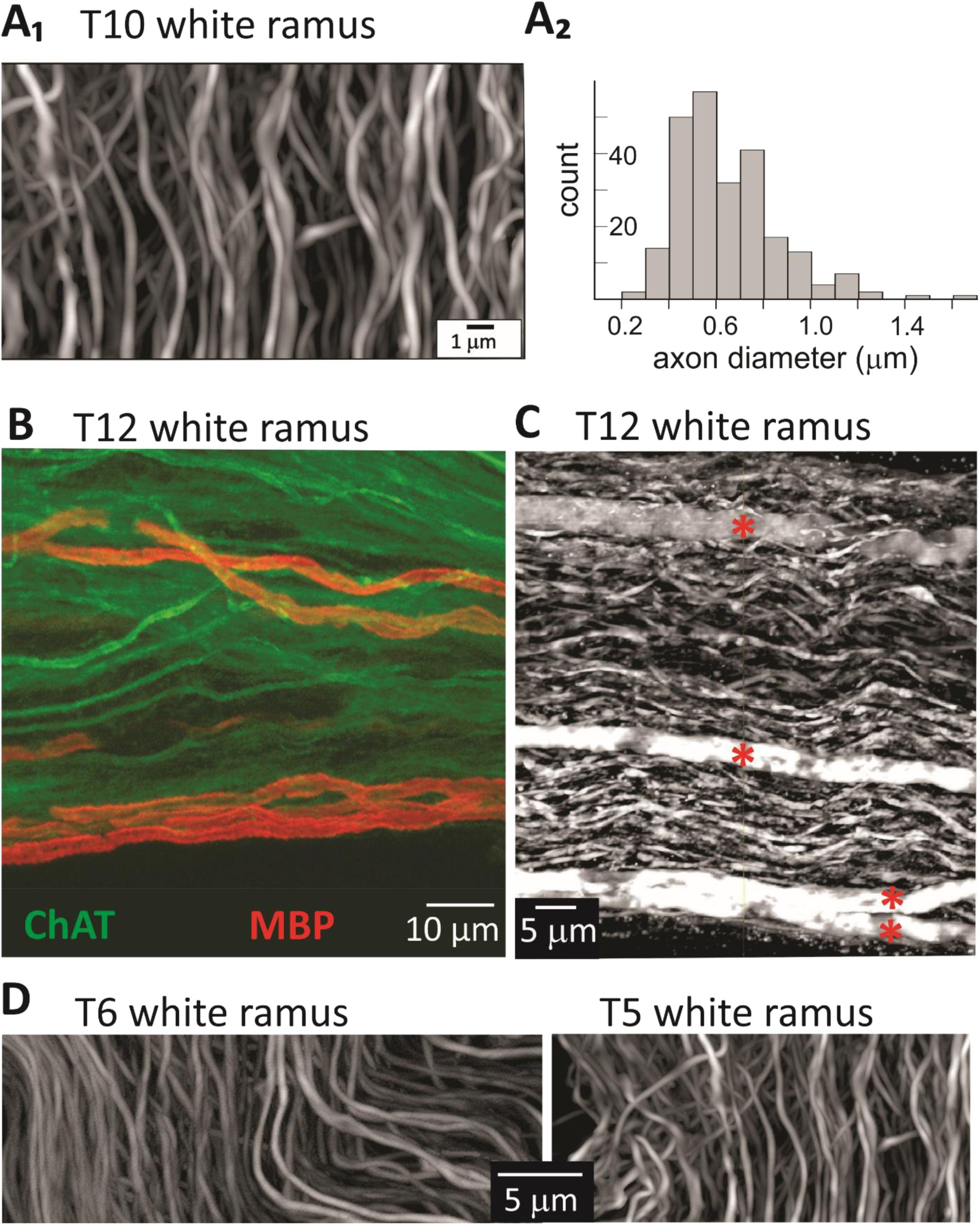
Thoracic SPN axons are predominantly unmyelinated. **[A]** Axon composition in T10 white ramus labeled with Neurobiotin. **[A_1_]** Image stacks of white rami were collected on a Nikon CSU-W1 SoRa spinning disk confocal microscope (60x objective, 4x magnification) and deconvolved for optical resolution of 0.112 µm. To facilitate unbiased analysis, axon counts and sizes were determined in FIJI and reported as the mode of >100 slices. **[A_2_]** Estimated axon count in this white ramus is 241. Mean=0.64±0.22μm(SD). **[B, C]** SPN axon composition in the T12 white ramus. **[B]** T12 white ramus in a ChAT-YFP mouse reveals both unmyelinated (displayed in green) and myelinated SPN axons within the white ramus, the latter being identified with immunolabeling for myelin basic protein (MBP; red). **[C]** T12 white ramus in ChAT-TdTomato mouse. Confocal scan of the white ramus, showing both myelinated (red asterisks) and unmyelinated axons. **[D]** T6 and T5 white rami are shown for comparison in other mice with Neurobiotin labeling from their corresponding ventral roots. Note axonal morphology in all white rami include varicosities with tortuous trajectories.

In comparison, caudally-projecting axons appear to have fewer axons entering the ganglion (T8) and had overall fewer axons exiting distally (**Fig. 1B_4_**). Instead, many axons bypassed the ganglion as unbranching axons projecting to the splanchnic nerve (**Fig. 1B1**).

Note that greater numbers of rostrally projecting axons were seen exiting than entering both the T6 and T5 ganglia (**Fig. 2A**), demonstrating that branching has occurred within these ganglia prior to projecting to additional ganglia as independent daughter branches. That axon counts were lower at more distant sites from Neurobiotin injection (**Fig. 2A**; see T4 and T3) is consistent with distance dependent loss of Neurobiotin label in many axons - indicating that branching is likely underestimated. Additionally, many axons may end blindly within the IGN.

We measured axon sizes in the IGN to determine whether axon diameters change with distance from the injection site. To compare axon diameter and numbers, we estimated values across 120 transverse consecutive optical slices in the IGNs across T7-T6, T6-T5, and T5-T4. Using this method, we observed increase in axon counts further from the diffusion site (**Fig. 2B**) but axon diameters in the IGN were comparable throughout at 0.36, 0.37, and 0.35µm at T7-T6, T6-T5, and T5-T4 IGNs, respectively (**Fig. 2C**). These values are comparable to the mean diameter of 0.4 µm from a prior study of superior cervical ganglion (Lewis and Burton, 1977). The distribution of axons the T7 injection site spreads out within interganglionic nerve at more distant sites presumably due to interspersed axons from other segments (**Fig. 2D**). Note axonal morphology in all the IGN includes varicosities with tortuous trajectories.

Consistent with anatomical projections patterns, SPN axons originating from various thoracic ventral roots exhibited patterns of both convergence and divergence across multiple ganglia (**Fig. 3A**). Notably, the largest CAPs generally occurred in the nearest ganglia. The T12 ganglion consistently received convergent input from all rostral segments up to T6 (n=4/4), while convergence from further rostral segments was not consistent (T5 – n=3/4; T3 – n=1/4). This distribution is consistent with that previous reported in guinea pig (Lichtman et al., 1980). Overall, these results demonstrate that SPN axons project to influence multiple segments and can establish divergent connections to multiple ganglia.

### Conduction variability studies

#### Evidence of spike conduction failures

Considering that axon diameters of SPN axons were ∼0.4 µm in the IGN and much smaller within the extensive branches within ganglia (Lewis and Burton, 1977), we sought evidence of conduction propagations failures, contrary to the previously assumed viewpoint of reliable conduction (McLachlan, 2003). We first sought to identify evidence of individual axons undergoing variation in recruitment consistent with conduction slowing and block (De Col et al., 2008; Pekala et al., 2016).

We recorded evoked responses in the caudal-end of the T11 ganglion (T11 IGN) following supramaximal stimulation of SPN axons in T7-T9 ventral roots (**Fig. 3B**). As expected, evoked responses were progressively smaller at stimulation sites from more distant spinal segments as less SPNs axons innervate distant ganglia (see also, **Fig. 5A_1_**). Importantly, we observed clear time-dependent evidence of variability in recruitment of SPN axons originating from all three segments (**Fig. 3B**). Presentation of the first 50 minutes of collected data as a raster plot clearly shows individual earlier-arriving events appearing and disappearing over time, with two emergent events undergoing progressive leftward shifts. Because reductions in propagation reliability are frequently associated with conduction slowing and eventual failure, the observed leftward shifts in latency and narrowing of temporal dispersion are consistent with improved propagation reliability (Debanne, 2004) (**Fig. 3C_1_**). The high variability in recruitment of the more numerous later-arriving events prevented identification of individual axon failures (blue shaded region) but clear time-dependent changes in recruitment can be seen when comparing population responses at different 5-min epochs (**Fig. 3C_2_**). Characterization of this variability required quantification as population changes as described below.

**Figure 5.**
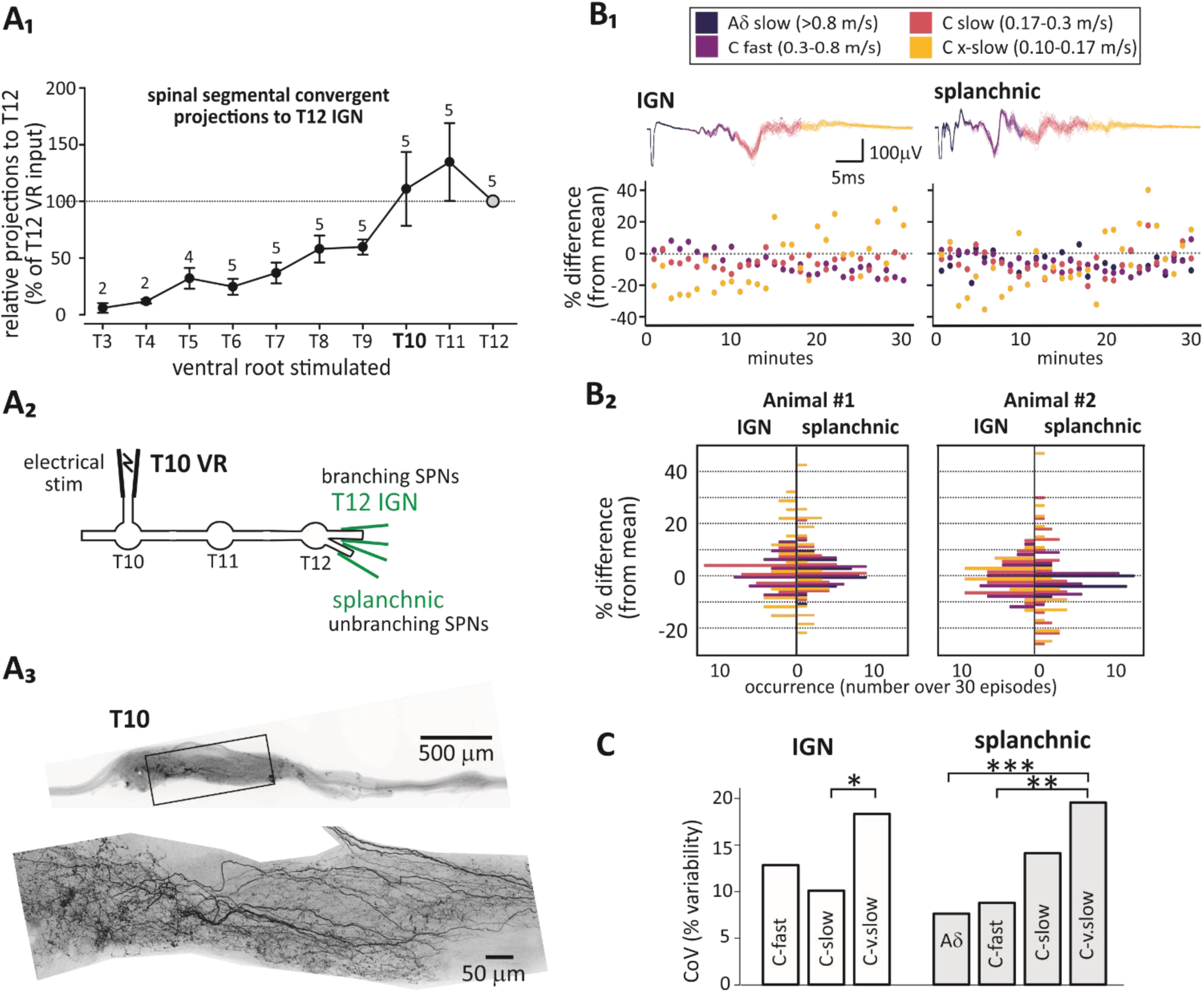
Spontaneous conduction block and emergence is seen in all fiber types. **[A_1_]** Relative multisegmental convergent projections to T12. Values represent the mean rectified integral population responses expressed as % of T12 amplitude. Sample sizes are in parentheses above each segment stimulated. **[A_2_]** Experimental setup showing SPN electrical stimulation in the T10 ventral root (VR) while recording evoked CAPs from the caudal T12 IGN and the splanchnic nerve. **[A_3_]** T10 SPN axonal labeling following Neurobiotin labeling of T10 VR. Note axonal branching within the caudal portion of the T10 ganglion prior to entry into the IGN. **[B_1_]** Showcase of CAPs recorded with electrical stimulation, divided into epochs based on conduction velocity. Superimposed episodes (10) illustrate the increased jitter as conduction velocities decrease. Aδ fibers, exclusive to the splanchnic nerve, are shown. Time-series plots from one animal demonstrate CAP rectified integral variations from the mean over 30 minutes. **[B_2_]** This panel presents histograms plotting the percentage change from the mean in two different animals, highlighting the variability across fiber types. Note that the extra-slow C fibers typically showing the largest percentage change from the mean. **[C]** Coefficient of variation (CoV) differs across fiber types. Mean CoVs show highest variance in the slowest axons (n=7; *p<0.05, **p<0.01, ***p<0.001).

### Comparing sensitivity to conduction failures across populations separated by conduction velocity

#### Identification of axon populations by conduction velocity

As all studies on the conduction velocities of SPN axonal populations were undertaken via recruitment at T10, we undertook high-resolution imaging at the T10 white commissure (ramus) following NB labeling. Total axon counts were 241 with mean diameter of 0.64±0.22μm (SD) (**Fig. 4A**). Axon diameter range was similar to that in the superior cervical ganglion (Lewis and Burton, 1977), and included a few axons with diameters typical of smaller diameter myelinated Aδ fibers. Previous reports have indicated that in the superior cervical ganglion (Lewis and Burton, 1977), about 99% of mouse paravertebral SPN axons are unmyelinated. In a separate mouse, co-labelling of cholinergic axons combined with immunostaining for myelin basic protein verified the presence of a small number of larger myelinated axons in the white commissure (**Fig. 4B,C**). These results are consistent with studies described below showing higher conduction velocities (CVs) in some axons consistent with slowly-conducting myelinated axons. NB labeling in other mice of the T6 and T5 white commissure show a similar axonal composition (**Fig. 4D**) to demonstrate that axonal morphology in all white rami include varicosities with tortuous trajectories.

As we were particularly interested in understanding conduction failures in the paravertebral ganglia, we first assessed convergent axonal contributions from various spinal segments. Relative amplitude contributions onto caudal T12 IGN from spinal segments T3-T12 are shown in (**Fig. 5A_1_**). Predominant contributions arise from T10-T12 with axons in T10 representing ∼19% of descending paravertebral SPNs projecting through the T12 ganglia.

All studies on the conduction velocities of SPN axonal populations were undertaken at following block of synaptic transmission with hexamethonium while independently recording from the IGN caudal to the T12 ganglia (i.e. across 3 ganglia) and the adjacent splanchnic nerve of the same length (see schematic **Fig. 5A_2_**). The splanchnic nerve is composed of unbranching SPN axons that project to prevertebral ganglia. As such it provides comparison to spike conduction properties of branching IGN axons. Anatomical examinations using Neurobiotin labeling of SPN axons from the T10 ventral root confirmed that SPN trajectories include branching axons across ganglia (**Fig. 5A_3_**).

The measured distance from the T10 ventral root to the splanchnic or IGN was determined to be ∼5mm. Conduction velocities (CV) for electrically stimulated SPNs were determined at 22°C (n=7). The CV range of the earliest and latest units arriving units in the CAP is provided in **Table 1**. The splanchnic nerve possessed fibers that conduct at faster velocities compared to those in the IGN but also contained the slower CV populations seen in the IGN.

**Table 1.** Conduction velocities as 22°C. Conduction velocity range of thoracic SPN axons in the interganglionic nerve (IGN) and splanchnic nerve (Spl) in the mouse at 22°C. SPNs were recruited with supramaximal stimulation of the T10 VR while recording from the caudal T12 IGN and the adjacent Spl nerve at equivalent distance from stimulation site (∼4.4mm).

| T10 VR stim (22°C) | Electrically Evoked |
| --- | --- |
| Interganglionic nerve | 0.11-0.84 m/s |
| Splanchnic nerve | 0.13-2.00 m/s |

First, we divided the evoked CAP into four conduction velocity ranges (**Fig. 5B**). These categories align with conduction velocities previously identified with specific afferent fiber types at 22C (Pinto et al., 2008). The fastest fibers likely correspond to lightly myelinated axons, possessing conduction velocities resembling those of Aδ fibers (>0.8 m/s). Notably, these fibers appear exclusively in splanchnic nerve recordings. The other three CV classifications fit within the C-fiber conduction velocity range characteristic of unmyelinated axons. To assess whether differences are seen within the C-fiber class, we divided the C-fiber component of the CAP into 3 epochs and labelled them as fast C-fibers (0.3-0.8m/s), slow C-fibers (0.17-0.3m/s), and extra-slow (x-slow) C-fibers (0.10-0.17m/s) (see **Fig. 5B**).

#### Evoked response fidelity is high in the fastest axons whereas the slowest axons have high response variability indicative of high rates of conduction failures

We compared variability in the rectified integral CAP amplitude evoked responses across these velocity ranges, gathered every minute over 30 minutes. We observed greater inter-episode time-dependent variability population recruitment in the later arriving fiber types (**Fig. 5B_1_**). The variability in fiber type responses between T12 IGN and splanchnic nerves is illustrated through histograms in two separate experiments, showcasing the diversity in percent change from the mean and the similarity in distribution across different animals (**Fig. 5B_2_**).

To assess how consistent variability in conduction was seen between mice and across axonal populations groups, we standardized dispersion by using the coefficient of variation (**CoV**) across fiber types in 7 mice (**Fig. 5C**). We observed a trend for increasing CoV % values correlating with decreasing conduction velocity in both splanchnic and IGN axons (ANOVA p<0.001 and p<0.05, respectively). Post-hoc analysis showed that x-slow C fibers had significantly higher CoV than Aδ (p<0.001) and C-fast fibers (p<0.005) within the splanchnic nerve, and slow C fibers in the IGN (p<0.05).

#### Activation history significantly reduces conduction

Activation history can alter preganglionic spike conduction properties (Zhang et al., 2017). This would have physiological relevance to the recruitment of postganglionic neurons. The effect of activation history on population recruitment was investigated by comparing evoked responses following delivery of stimuli at 1Hz, 5Hz and 10 Hz interpulse intervals. The 1 Hz test involved single pre-train stimulus delivered 1-second before delivery of 5 or 10Hz trains (2-second duration). As pre-pulses were delivered one second before both frequency trains, this 1 Hz data was pooled (n=7) and showed that, after a single pulse, evoked responses were significantly depressed in IGN (to 85.1±4.5% of control; p=0.015; paired t-test) but not in splanchnic (to 98.9±4.0% of control; p=0.8; paired t-test).

Additional 5Hz and 10 Hz results are presented in Figure 6 and Table 2. As shown, 5Hz or 10Hz frequency-dependent responses progressively depress in amplitude, then largely stabilize by the 5th-6th pulse (**Fig. 6A**). Over 9 consecutive stimuli, there was an overall significant effect of activation history on reduction in response magnitude in IGN at both frequencies (p<0.05; ANOVA on ranks). Moreover, response reduction was significantly greater at 10 vs. 5 Hz (p<0.05; t-test). In comparison, numerical reduction in magnitude was smaller in response magnitude in the splanchnic nerve but still significant at 5Hz (p<0.05; ANOVA on ranks).

**Figure 6.**
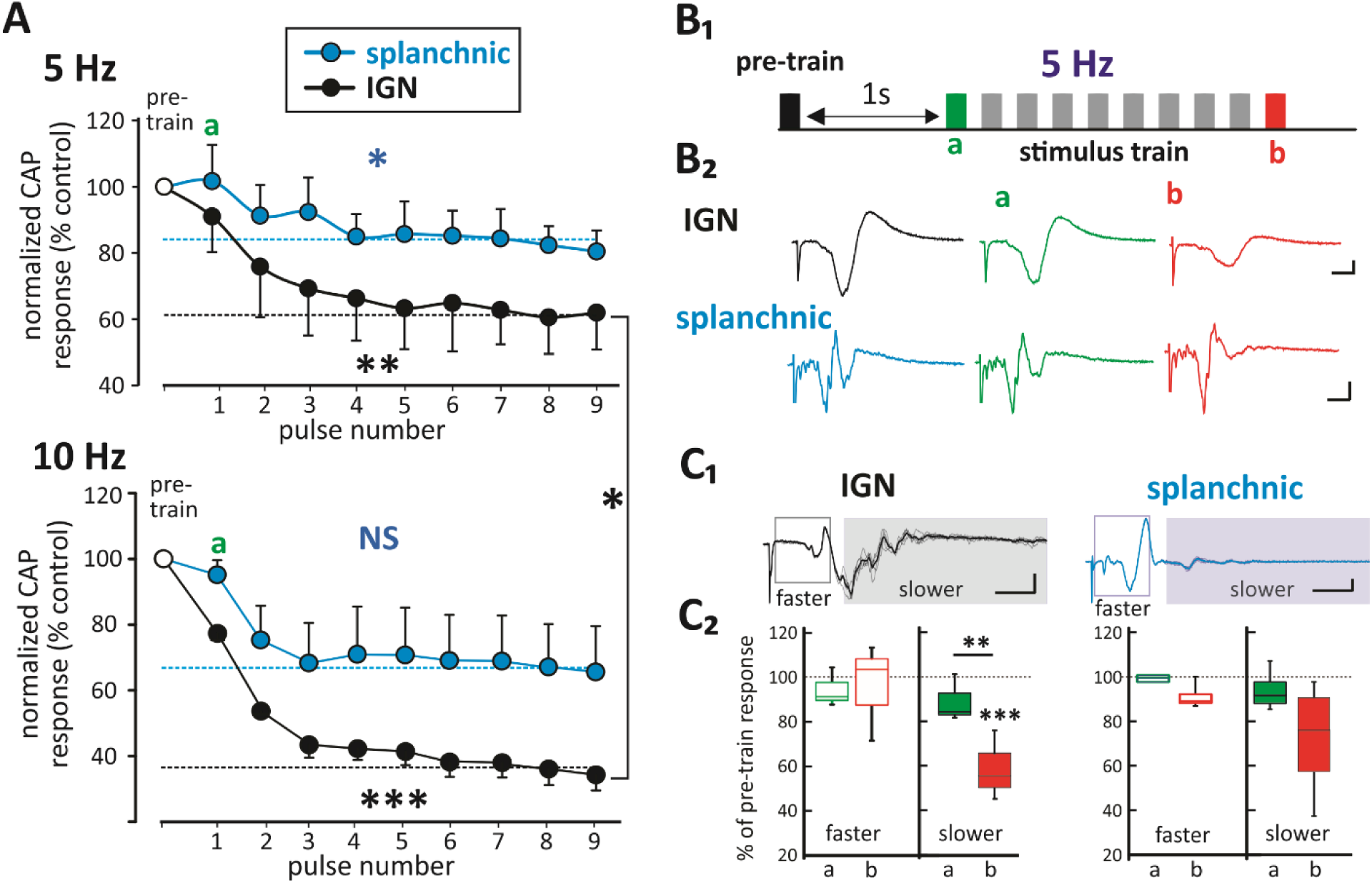
Slower-conducting IGN axons undergo preferential depression to stimulus trains. **[A]** Effect of pulse number on change in evoked CAP response amplitude during 10 and 5Hz stimulation. Percentage changes in CAP responses relative to the pre-train pulse are depicted Typically, maximum depression response was seen by 6th pulse (10 Hz [n=3]; 5 Hz [n=4]). There was an overall significant effect of activation history between the 1^st^– 9^th^ pulse on reduction in response magnitude in IGN at both frequencies (** and ***; RM ANOVA on ranks), and the reduction was significantly greater at 10 vs. 5 Hz (* at right; t-test). SPL response at 5Hz but not 10Hz also underwent significant effect of activation history (RM ANOVA on ranks). **[B]** Experimental design. **[B_1_]** Experimental design for 5Hz, 2 sec stimulus train initiated 1 sec after the pre-train response. Quantified are responses at train start (**a**) and train end (**b**). **[B_2_]** Average responses of first and last response in 5 Hz train in both the IGN and splanchnic nerve in the same mouse. **[C]** Comparing fast and slow responses in the 5Hz train (for 10 Hz see Table 2). **[C_1_]** Example of 5 raw and superimposed average value of evoked responses separated into faster and slower components. **[C_2_]** Rectified-integral values of both faster- and slower-conducting components expressed a percentage of the corresponding pre-train pulse values. Frequency-dependent depression was seen in the slower-conducting axons in the IGN only (paired t-test; epoch (**b**)). *, p<0.05; **p<0.01; ***p<0.001. Scale bars are 50 μV, 10 ms.

**Table 2.** Effects of frequency trains on response magnitude relative to prepulse delivered 1 sec prior. Comparing frequency- and time-dependent changes in evoked responses to 2 second trains at 5Hz (n=4) & 10Hz (n=3). Responses are separated into fast (0.3-2.2 m/s) and slow (0.06-0.2 m/s) SPNs in the IGN and Spl nerves. Epochs **a** & **b** provide the amplitude responses of the 1st and last pulse in the train, respectively. Amplitudes are relative to a pre-train evoked response delivered 1 sec prior to train onset. Values represent the mean % of pre-train response ± SEM with significance reported as; ** p<0.01, and *** p<0.001.

| epoch | Fast SPNs (0.3-2.2 m/s) |  | Slow SPNs (0.06-0.2 m/s) |  |
| --- | --- | --- | --- | --- |
|  | 5Hz | 10Hz | 5Hz | 10Hz |
| **IGN** |  |  |  |  |
| **a** (1<sup>st</sup>) | 94.4 ± 8.5 | 92.5 ± 14.6 | 89.1 ± 10.5 | 85.0 ± 12.9 |
| **b** (last) | 96.0 ± 21.8 | 188.6 ± 90.6 | 58.9 ± 15.9*** | 25.4 ± 11.6*** |
| **Spl.** |  |  |  |  |
| **a** (1<sup>st</sup>) | 97.8 ± 3.52 | 100.4 ± 6.7 | 92.9 ± 9.5 | 67.5 ± 45.8 |
| **b** (last) | 90.2 ± 6.0 | 91.6 ± 10.9 | 70.7 ± 27.1 | 51.1 ± 27.7 |

Quantification of activity-dependent changes in the CAP at 22C were based on the experimental design shown (**Fig. 6B**). A rest period of 90 seconds between trials ensured recovery to baseline response. As shown, we compared the pre-train CAP, evoked one second before the sequence began, to the 1^st^ response (1Hz; epoch [a]) and last response within the stimulus train (epoch [b]). Example average electrically-evoked responses are illustrated in **Fig. 6B_2_**, respectively. All signals were high pass filtered at a frequency of 25Hz.

In contrast to the methodology above, where we divided our analysis into four CV classes, we chose a different approach for these frequency-dependent studies. This decision was made to accommodate potential changes in axonal conduction velocity that could arise from varying frequencies. Consequently, evoked responses for each epoch were divided into faster- (0.3-2.2 m/s) and slower-conducting (0.06-0.2 m/s) components (**Fig. 6C**).

Frequency-dependent decreases were preferential to slower-conducting events in the IGN at both frequencies (**Fig. 6C_2_**; **Table 2**). The average response of slow IGN axons at the end of the 5 and 10 Hz trains (epoch [b]) dropped significantly to 59±16% and 25±12% of the pre-train response, respectively (p<0.001 for both; **Table 2**) with the reduction at 10Hz being significantly lower than seen at 5 Hz (t-test; p<0.05).

## DISCUSSION

### Summary of principal findings

Sympathetic preganglionic neurons (SPNs) represent the final common pathway for central sympathetic output, yet the reliability of spike propagation along their axons has largely been assumed rather than directly examined. Classical descriptions of autonomic transmission assume that sympathetic signals propagate faithfully along SPN projections (McLachlan, 2003), and emphasize synaptic integration within the spinal cord and ganglia as the primary determinants of sympathetic drive (Janig, 2022). As shown in the summary **Figure 7**, the principal conceptual advance of this work challenges the prevailing view by identifying axon conduction reliability as a previously underappreciated presynaptic control point, indicating that variability in spike propagation within the preganglionic axon population itself can influence the number of postganglionic neurons recruited by a given central command. Such axonal mechanisms therefore represent an additional level of regulation that complement synaptic and network-level control of autonomic function.

**Figure 7.**
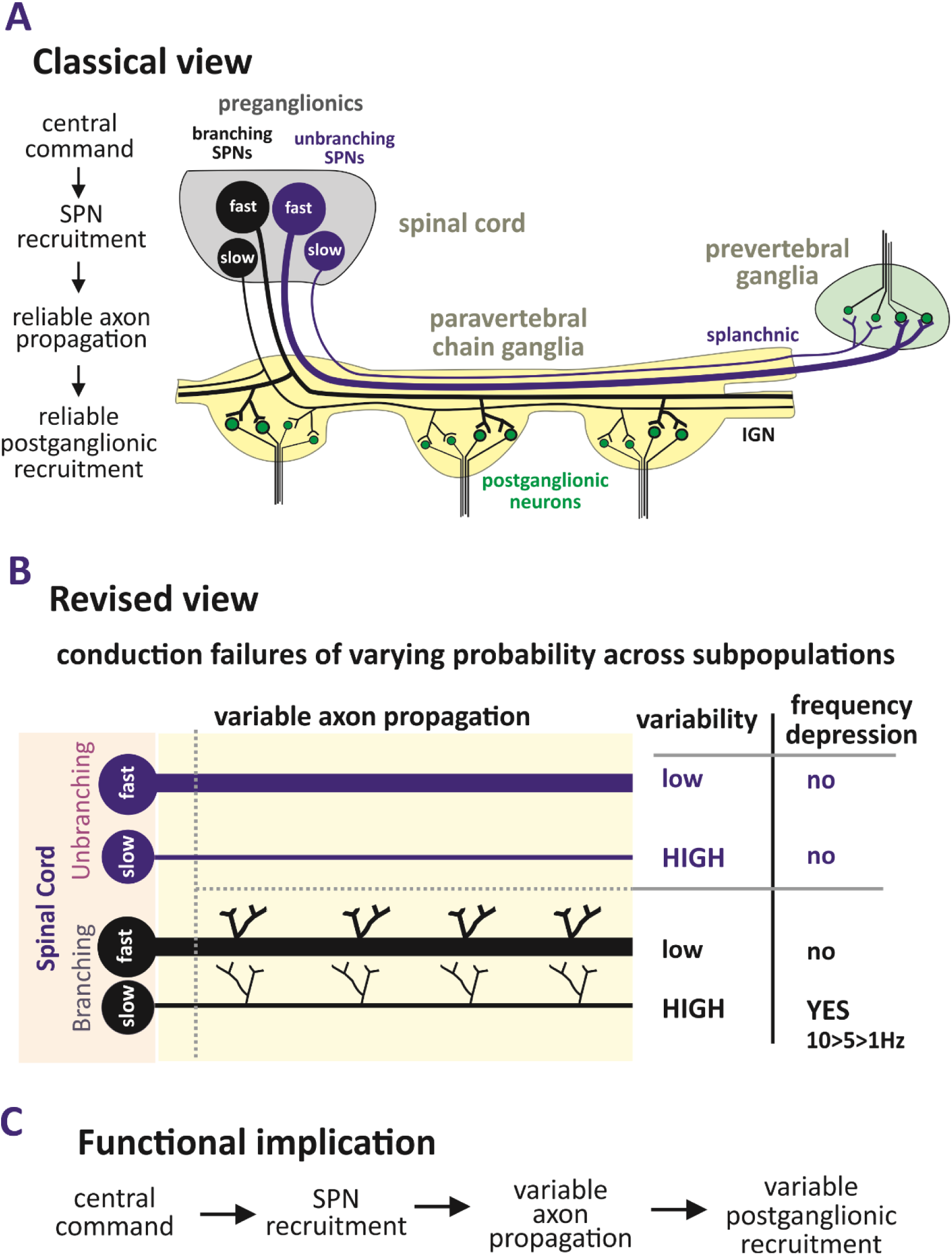
Axonal conduction failures modulate sympathetic output. **[A]** Summary anatomical organization and associated classical view in which spikes propagate reliably along divergent sympathetic preganglionic neuron (SPN) axons, including across multiple branch points, for reliable recruitment of postganglionic neurons in sympathetic ganglia. **[B]** Revised view synthesizes the anatomical reality of very small unmyelinated axons with irregular morphology and branching populations with the experimental observation of probabilistic conduction failures across repeated trials, particularly among slow-conducting unmyelinated axons. Slow-conducting branching axons within the paravertebral sympathetic chain are uniquely sensitive to activation history-dependent conduction failures that increase with increased frequency. **[C]** Proposed functional model: conduction variability acts as a presynaptic gain control node shaping sympathetic population output. Variability is probabilistic and can modulate a given central sympathetic command.

The present study demonstrates substantial intertrial variability in SPN evoked responses consistent with frequent failures of spike propagation. Variability was not uniformly distributed across axonal populations. Moreover, variability was not observed uniformly across axonal populations. It was most pronounced in the slowest-conducting axons with preferential increased failures during repetitive activation in branching axons. If high rates of propagation failure are characteristic of SPN populations, it suggests sympathetic output is driven by probabilistic rather than deterministic features of population encoding.

### Extensive divergence of SPN axons across sympathetic chain ganglia

Using an ex vivo preparation preserving multisegmental sympathetic chain connectivity, we undertook Neurobiotin anterograde labelling of SPN axons from individual thoracic ventral roots. Results demonstrated that SPN axons exhibit extensive rostrocaudal projecting across multiple thoracic ganglia and consisted predominantly of small diameter unmyelinated fibers. Consistent with previous reports in superior cervical ganglia (Lewis and Burton, 1977), SPNs in the mouse paravertebral thoracic chain were shown to be overwhelmingly small-diameter, with only a small number of larger diameter myelinated axons present. This was consistent with the predominantly unmyelinated fiber conduction velocity measures obtained in SPN axons recorded in splanchnic (unbranching) and IGN (including branching) axons. While the splanchnic nerve contained faster conducting myelinated axons both IGN and splanchnic had a comparable range of slower conducting axons whose velocities were consistent with those of unmyelinated axons (Pinto et al., 2010).

Many axons bifurcated within the white ramus to generate both rostral and caudal projections, allowing a single SPN to influence postganglionic neurons distributed across several segments. These observations are consistent with classic studies demonstrating multisegmental divergence of SPN projections in guinea pig and rat preparations (Blackman and Purves, 1969; Nja and Purves, 1977; Lichtman et al., 1980; Forehand et al., 1994). Divergent projections enable substantial amplification of central sympathetic commands, with each SPN capable of influencing many postganglionic neurons (Purves et al., 1986).

Our anatomical labeling of T7 SPN axons revealed intricate branching patterns within the thoracic chain ganglia with notable ‘T’ shaped branch points bifurcating within the T7 ganglia to enable rostral and caudal projections. While T-shaped trajectories were observed, complete reconstruction of individual axons was not feasible, and the prevalence of true bifurcations remains uncertain. A prior study on projections from in rat (embryonic and adult) and chicken reported only a small % of axons bifurcate while the other axons only project rostrally or caudally (Forehand et al., 1994). As the Forehand et al (1994) study used different fluorescent retrograde dyes in rostral and caudal IGN across ganglia, is possible these dyes had limited anterograde transport across rostrocaudal bifurcations (Lanciego and Wouterlood, 2020).

Analysis of rostral projections show that many axons branched to form synapses at T6 and T5 before continuing rostrally with a significant decrease in axon number observed rostral to the T4 ganglion. Axon counts exiting were greater than those entering T6 and T5 consistent with interganglionic branching with daughter branches reentering the IGN. However, there was no substantial change in axon diameter along the measured interganglionic pathway.

Our electrophysiological studies revealed that SPN axons, originating from mid- to caudal-thoracic spinal segments, demonstrate significant multisegmental convergence onto recorded T12 IGN. A similar multisegmental convergence was previously reported in guinea pig (Lichtman et al., 1980). Notably, the T12 ganglion showed convergence from segments rostral to T12 up to T6 consistently, but this pattern varied above T6. Our finding that SPN axons arise from multiple spinal segments with the largest CAPs typically observed in the nearest ganglia is consistent with previous reports in guinea pig thoracic ganglia (Lichtman et al., 1980).

### Small axon diameter and susceptibility to conduction variability

Consistent with previous reports, the vast majority of SPN axons observed in the present study were unmyelinated with diameters generally well below 1 μm (Lewis and Burton, 1977). Small-diameter axons are predicted to exhibit reduced safety factors for spike propagation due to high axial resistance and large surface-to-volume ratios (Debanne, 2004; Debanne et al., 2011). We undertook studies at room temperature (22C) to align observed responses in relation to the only other study that assessed actions across thoracic ganglia at 22C (Lichtman et al., 1980). Relative to physiological temperatures, reduced temperature would be expected to broaden action potentials and increase safety margins for propagation (Hodgkin and Katz, 1949), so it was unexpected to discover that, even at 22C, slower conducting axons demonstrated high variability in response conduction. That unbranching splanchnic SPNs showed a similar degree of variability in response in slow conducting axons as branching axons in the IGN suggests that the diameter of the smallest axons may be a dominant determinant of propagation reliability. Similar diameter-dependent conduction variability has been observed in other populations of unmyelinated axons (Debanne et al., 2011). Importantly, at physiological temperatures, conduction failures would be expected to increase due to effects on membrane resistance (Maingret et al., 2000) and spike shape (Hodgkin and Katz, 1949). This is explored separately in another study(Halder and Hochman, 2026).

### Frequency-dependent modulation of conduction reliability

Our exploration into the frequency-dependent transmission in SPN axons is grounded in the context of existing literature. SPNs typically discharge at low rates, often near or below 1 Hz, although bursts reaching approximately 10 Hz and higher have been reported (McLachlan, 2003; Janig, 2022) though higher firing frequencies has also been reported (Janig et al., 1982). We examined the effects of two stimuli at 1 Hz and during 5 and 10Hz stimulus trains on SPN amplitude changes. Experiments using repetitive stimulation exhibited depression during stimulus trains, particularly in branching interganglionic nerves. In contrast, faster-conducting axons remained relatively stable across stimulation frequencies. Frequency-dependent depression may arise from several mechanisms including sodium channel inactivation, extracellular K^+^ accumulation, or Na⁺/K⁺ pump–mediated hyperpolarization (Dobretsov and Stimers, 2005; Zhang and Sillar, 2012). Such mechanisms could transiently reduce the safety factor for spike propagation in small axons following periods of activity.

### Branch points as potential sites of conduction failure

Axonal branching introduces geometric discontinuities that can reduce the safety factor for spike propagation. Theoretical and experimental studies have shown that propagation across branch points may fail when the impedance mismatch between parent and daughter axons becomes sufficiently large (Goldstein and Rall, 1974; Levy, 1980; Debanne, 2004; Hari et al., 2022). These mechanisms are particularly relevant in systems with extensive axonal arborization. As stated, while evidence of variability was observed in both branching and nonbranching pathways, frequency-dependent failures were preferential to and profound in branching IGN axons. The sympathetic chain provides numerous opportunities for such branch-point effects because SPN axons form both bifurcating and collateral branches along their rostrocaudal trajectory.

### Implications for sympathetic population coding

The sympathetic nervous system operates primarily through changes in the population activity of SPNs rather than through precisely timed single-axon signaling. Under such conditions, probabilistic spike propagation in small-diameter axons may serve as a mechanism that modulates the effective size of the recruited postganglionic population. This is consistent with observations that recruitment of individual postganglionic neurons is also probabilistic (Macefield and Wallin, 1999, 2018). Small axons also represent an energetically efficient strategy for maintaining large numbers of neurons within autonomic circuits (Perge et al., 2012). This metabolic efficiency likely comes at the expense of reduced conduction safety factors. The resulting conduction variability may therefore represent a trade-off between energetic efficiency and signal reliability in autonomic systems.

### Limitations

Several limitations should be acknowledged. First, conclusions regarding spike propagation failures are inferred from population CAP measurements rather than direct recordings from identified axons. Consequently, altered excitability, recruitment thresholds, or temporal dispersion could contribute to observed changes. Second, axonal subtype classification relied on conduction velocity rather than direct morphological reconstruction. Finally, experiments were performed *ex vivo* under controlled conditions and may not fully capture the influence of ongoing neuromodulatory states present *in vivo*.

### Perspective and impact

Classical estimates of sympathetic amplification are based primarily on anatomical ratios between pre- and postganglionic neurons (Wang et al., 1995). However, our observations suggest that propagation reliability may substantially influence the effective recruitment of downstream populations, particularly in the slowest conducting presumably more highly branched axonal projections. The specific mechanisms underpinning the observed low safety factor in conduction, including in relation to different SPN types, should provide additional insight into physiological relevance and potential dysfunction in disease.

## Acknowledgements

This research was funded by NIH NS121850, NS102871

